# Ephrin Signaling Patterns Sensory Neurons During Tissue Homeostasis in Planarians

**DOI:** 10.64898/2026.08.11.744258

**Authors:** Mohammad A. Auwal, Sarah E. Warner, Allisyn Marks, Ryan A. McCubbin, Arianna L. Farrar, Jennifer M. Severance, Christian Torres, Kelly G. Ross, Ricardo M. Zayas

**Author notes:** **Correspondence:** Ricardo M. Zayas. Equal authorship (Mohammad A. Auwal, Sarah E. Warner).

## Abstract

Eph and ephrin genes encode receptor-ligand pairs that mediate contact-dependent cell signaling and are essential for nervous system development. However, less is known about the role of Ephrin signaling during adult tissue homeostasis and regeneration. Here, we investigated the role of Ephrin signaling in neural patterning in the planarian *Schmidtea mediterranea*. We discovered that RNAi against the Eph receptor *EphR1* led to striking ectopic expression of the mechanosensory neuron markers *pkd1L-2* and *hmcn-1-L,* without obvious disruption of the overall architecture of the central nervous system. To investigate the basis of this phenotype, we identified additional Eph receptor homologs and four putative ephrin ligands and assessed their function. An RNAi screen revealed that *ephrin*-*1* phenocopies the defects of *EphR1* RNAi. Temporal analyses of *EphR1* and *ephrin-1* inhibition revealed a progressive increase in *pkd1L-2^+^* and *hmcn-1-L^+^* cells, indicating an unappreciated role for Ephrin signaling in regulating neural patterning and cell number during adult tissue homeostasis. Together, these findings provide a framework for dissecting Ephrin-dependent mechanisms in adult tissue maintenance and regeneration.

## Introduction

The nervous system comprises extensive networks of neurons that relay information throughout the body. These cells acquire distinct identities based on their spatial position through developmental patterning, a process regulated by signaling gradients and guidance molecules (Kandel et al., 2014). In developmental neurobiology, guidance molecules such as Slits, Netrins, Semaphorins, and Ephrins direct cell migration and guide axons along attractive or repulsive substrates (Aberle, 2019). Ephrins are implicated in many processes, including cell repulsion, cell-cell adhesion, axon guidance, cell migration, cell proliferation and differentiation, and tissue boundary formation (Kania and Klein, 2016; Klein, 2012; Lisabeth et al., 2013). Their function is evolutionarily conserved across metazoans, and distant homology has even been identified in choanoflagellates (Arcas et al., 2020). In the ascidian *Ciona intestinalis*, Ephrins underlie cell fate in several lineages via contact-dependent signaling, and Ephrin signaling contributes to proper progenitor population size by increasing or decreasing cell proliferation (Wilkinson, 2014). In mammals, Ephrin signaling plays an important role in neurogenesis and regeneration after nervous system injury. The vast majority of neurons in humans are generated prior to birth, and adult neurogenesis is restricted to the subventricular zone of the lateral ventricles and the subgranular zone of the hippocampus (Jiao et al., 2008). Ephrins are expressed within these regions and throughout the CNS (Laussu et al., 2014). In many cases, Ephrins act as negative regulators of adult neurogenesis (Jiao et al., 2008), and in response to CNS injury, many Ephrins are upregulated and often act to regulate axonal regeneration and glial scar formation (Goldshmit et al., 2004; Yang et al., 2018).

The planarian S*chmidtea mediterranea* is an excellent model organism for studying neurogenesis and neuronal patterning (Ross et al., 2017). S*. mediterranea* are capable of essentially unlimited regeneration and can restore all tissue types and their functions from even small tissue fragments (Ivankovic et al., 2019; Reddien, 2018; Reddien and Sánchez Alvarado, 2004). This extraordinary regenerative capacity is due to an abundant population of pluripotent stem cells, called neoblasts, that remain active throughout the lifespan of planarians (Newmark and Sánchez Alvarado, 2002). Upon injury, these specialized cells proliferate and differentiate to replace any missing tissues, including neurons (Reddien, 2018; Ross et al., 2017). Studying neoblasts and their progeny can help elucidate mechanisms that coordinate processes required for development and regeneration, such as cell proliferation, differentiation, boundary formation, and assimilation into existing tissues. Although the planarian body plan is simple, the nervous system is relatively complex: numerous distinct neuronal populations are specified and patterned by highly conserved neural transcription factors and guidance molecules (Cebrià, 2007; Reddien, 2022; Ross et al., 2017).

Previous work from our laboratory revealed that the SoxB1 gene *soxB1-2* is critical for the regeneration and maintenance of the planarian mechanosensory system (Ross et al., 2018). Since this finding, we have uncovered a conserved regulatory network for the differentiation of a subset of ciliated mechanosensory neurons: *soxB1-2*^+^ neuro-epidermal progenitors commit to a *pou4-2^+^* neural lineage, differentiating into *polycystic kidney disease-1-like-2* (*pkd1L-2^+^*) or *hemicentin-1-like* (*hmcn-1-L^+^*) ciliated mechanosensory neurons. A screen for genes differentially expressed in *pou4-2^+^*cells identified an Ephrin Receptor 1 (*Smed-EphR1*) homolog as a candidate gene involved in mechanosensory organ patterning (McCubbin et al., 2025). Consistent with preliminary results from our laboratory (McCubbin, 2022), this study implicated *Smed-EphR1* as an essential regulator of mechanosensory neuron patterning. Inhibition of *EphR1* results in ectopic expression of *pou4-2^+^, pkd1L-2^+^,* and *hmcn-1-L^+^*ciliated mechanosensory neurons. Knockdown of *Smed-EphR1* also results in ectopic expression of *nkx2-4^+^*, *scratch^+^*, and *cav-1^+^* neural cells, suggesting that EphR1 plays a role in the patterning of neuron populations that might not be directly regulated by *pou4-2*. To uncover the mechanism underlying the *Smed-EphR1* RNAi phenotype, we identified additional Eph receptor and putative ephrin ligand homologs and assessed their expression and function. We found that *Smed-ephrin-1* is expressed in a pattern juxtaposed to *Smed-EphR1*, and *Smed-ephrin-1* RNAi phenocopies the *Smed-EphR1* RNAi patterning defects. To determine whether *EphR1-Ephrin-1* signaling regulates spatial patterning alone or also influences cell number, we performed temporal analyses while simultaneously inhibiting *EphR1* and *ephrin-1*. These experiments revealed a significant increase in the total numbers of differentiated *pkd1L-2^+^* and *hmcn-1-L^+^* cells, suggesting a role for *EphR1* in the feedback control of mechanosensory neuron neurogenesis.

## Results and Discussion

### *Smed-EphR1* is required for normal patterning of *pou4-2*-regulated ciliated stripe neurons

*In situ* hybridization experiments corroborated that *Smed-EphR1* (hereafter *EphR1*) was expressed in the head tip, the dorsal ciliated stripe, and the peripheral ciliated stripes (Fig. 1A), as shown in McCubbin et al. (2025). This stereotypical expression pattern is commonly observed in sensory neuron populations, including those involved in mechanosensation (McCubbin et al., 2025; Ross et al., 2024; Ross et al., 2018). Our previous work showed that inhibition of *soxB1-2* and *pou4-2* resulted in decreased expression of *EphR1* in the dorsal ciliated stripe and peripheral ciliated stripes, whereas expression in a population of cells at the head tip remained unaffected, indicating that *EphR1* signaling in most cells is downstream of *soxB1-2* and *pou4-2* (McCubbin et al., 2025). The overlapping expression patterns of *EphR1* and *pou4-2*-regulated ciliated stripe neurons suggested to us that *EphR1* might play a role in targeting this cell population to the appropriate anatomical locations.

**Figure 1.**
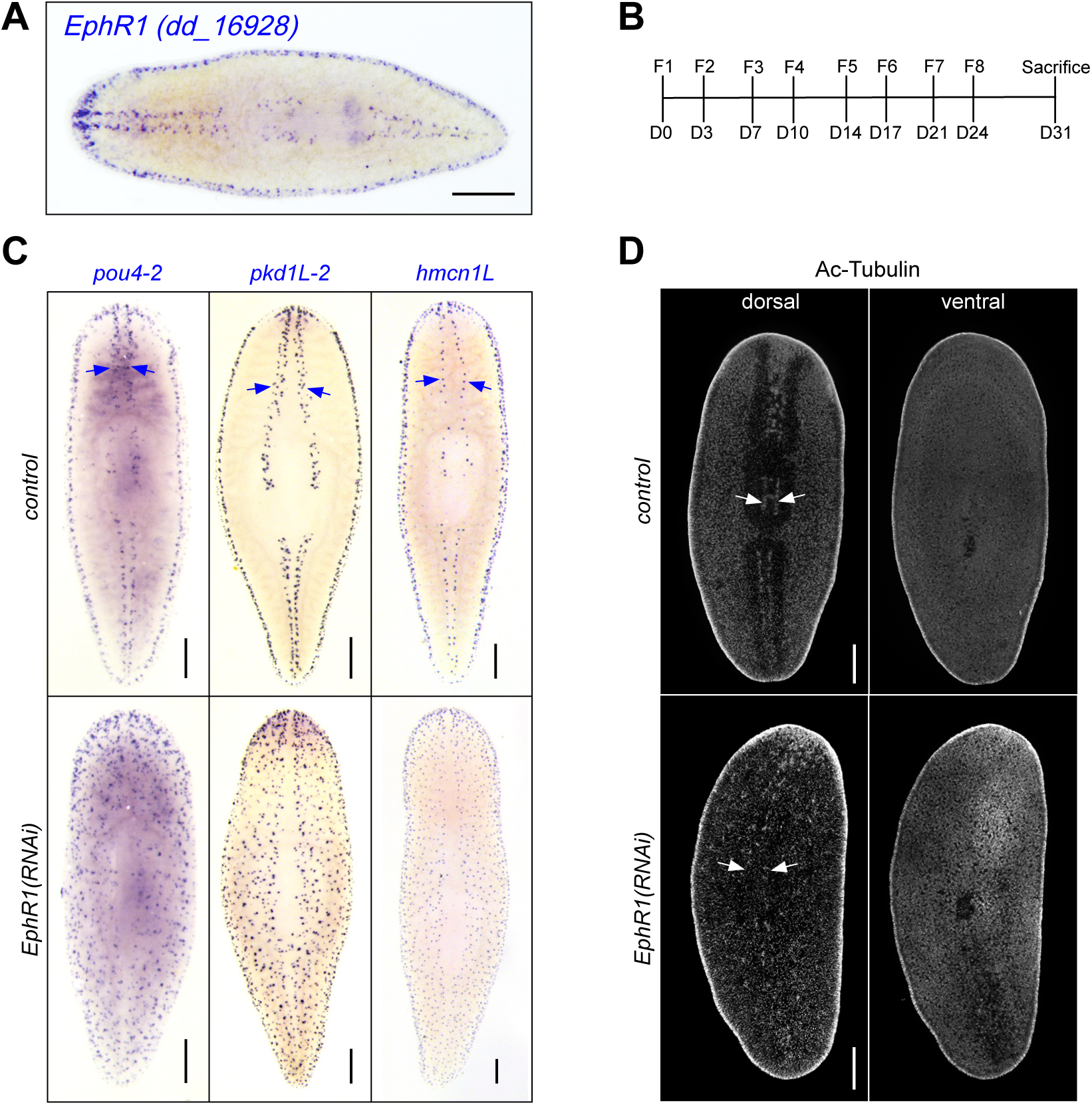
*EphR1* is expressed in the ciliated stripe and regulates its expression patterns. A) WISH analysis showed that *EphR1* is expressed along the dorsal head tip, the dorsal ciliated stripe, and the peripheral ciliated stripes. B) Knockdown of *EphR1* resulted in ectopic expression of *pou4-2^+^, pkd1L-2^+,^* and *hmcn-1-L^+^* cells along the dorsal ciliated stripe (blue arrows). C) *EphR1(RNAi)* animals showed disrupted dorsal ciliated patterning. Scale bars = 200 µm. Experiments were replicated 3 times, with 5 worms per condition and 1 representative worm imaged per condition.

To test our hypothesis, we knocked down *EphR1* using RNAi. We found that after a 31-day RNAi regimen consisting of eight dsRNA feedings over four weeks (Fig. 1B), *EphR1*(*RNAi*) animals displayed ectopic expression of *pou4-2^+^, hmcn-1-L^+^,* and *pkd1L-2^+^* cells (Fig 1C), and the dorsal ciliated stripe was lost. The cells marked by these genes were no longer confined to their normal boundaries and instead appeared scattered across the dorsal side of the body, with only the midline, medial to the stereotypical location of the dorsal ciliated stripe, remaining void of expression. This phenotype suggests that *EphR1* does not specify mechanosensory fate *per se*, but instead restricts where these neurons differentiate or persist within the peripheral nervous system architecture. Eph-ephrin signaling is well suited to this role because it provides contact-dependent, short-range cues that sharpen tissue boundaries and cellular distributions in other organisms (Kania and Klein, 2016). Our results extend these boundary-like functions to adult sensory neuron patterning during planarian homeostasis.

### *EphR1* regulates a specific subset of peripheral neurons in the planarian nervous system

Given the robust lateral expansion of mechanosensory neurons across the dorsal side of the planarian body, we wanted to examine whether EphR1 signaling broadly regulates planarian tissue patterning. First, we tested whether EphR1 signaling is necessary for the correct patterning of mechanosensory neurons regulated by POU4-2 by examining the expression of *scratch* and *calm2* after *EphR1* knockdown. Previous work determined that *scratch* and *calm2* are co-expressed with *pou4-2* (King et al., 2024; McCubbin, 2022)*. EphR1* RNAi resulted in ectopic expression of *scratch*^+^ and *calm2*^+^ neurons (Supplementary Fig. 1A), further confirming that the patterning of *pou4-2*-regulated ciliated neurons is regulated by *EphR1*.

We next examined *EphR1*(*RNAi*) planarians for changes in CNS patterning by assessing Synapsin expression after RNAi. Anti-Synapsin labels synapses and can be used to visualize the overall structure, with major labeling in the cephalic ganglia, ventral nerve cords, and pharyngeal nerve net (Cebrià, 2008). Knockdown of *EphR1* did not result in overt patterning defects in the cephalic ganglia, ventral nerve cords, or pharynx (Supplementary Fig. 1B). The preservation of major CNS structures argues against a broad requirement for *EphR1* in neural maintenance and instead supports a selective role in the organization of peripheral sensory neurons.

Thus far, we have evidence that EphR1 signaling regulates the patterning of *pou4-2^+^* mechanosensory cells without broad nervous system patterning defects. We next sought to determine whether other genes expressed in the mechanosensory organ but not under the *pou4-2* gene regulatory network would also be affected by loss of *EphR1* expression. We focused on two genes, *nkx2-4* (*dd_*33456) and *cav-1 (dd_8555*), that share expression patterns with *EphR1^+^* cells but function independently of *pou4-2* (McCubbin, 2022; Wang, 2019). WISH analysis revealed their expression within the mechanosensory neuron regions. Knockdown of *EphR1* resulted in ectopic expansion of cells marked by these genes (Supplementary Fig. 1C). The observed phenotype could reflect a direct requirement for EphR1 in these additional ciliated cell populations or an indirect consequence of altered signaling from mispatterned mechanosensory neurons. Distinguishing between these possibilities will require cell-type-specific expression and co-expression analyses. Thus, an important open question is whether these additional ectopic populations reflect direct *EphR1* expression within multiple peripheral neuron classes or indirect effects on neighboring cells within the ciliated stripe environment. Distinguishing these models will require co-expression analyses.

### Screening additional Ephrin signaling genes to identify potential *EphR1* functional partners

*EphR1* is not the only Ephrin signaling molecule encoded in the planarian genome. We searched the planarian transcriptome and predicted proteome and identified seven additional candidate Ephrin signaling genes (see Methods; Supplementary Table 3). Phylogenetic analysis resolved relationships among the EphR and ephrin homologs: three additional Eph receptors, which we named *EphR2*, *EphR3,* and *EphR4*, and four candidate ephrin ligand-encoding genes, which we named *ephrin-1, ephrin-2, ephrin-3*, and *ephrin-4* (Supplementary Figs. 2-3). We identified the *EphR4* and *ephrin-4* genes while drafting this manuscript by searching the predicted proteome database (see Methods); analysis of those genes is in progress. In the interim, WISH analysis of all other Ephrin signaling genes revealed their expression patterns in the planarian body (Supplementary Fig. 4). *EphR2* expression was detected in the brain, ventral nerve cords, pharynx, and along the body periphery (blue arrow in *EphR2,* Supplementary Fig. 4). We observed *EphR3* expression at the anterior and posterior poles (blue arrow marks the anterior pole in *EphR3*, Supplementary Fig. 4), as well as along the body periphery. The observation of expression at the poles raises the intriguing possibility of an interaction between *EphR3* and polarity-control genes in the planarian, such as *beta-catenin*, *Wnt*, and *hedgehog* (Petersen and Reddien, 2009; Yazawa et al., 2009)*. ephrin-1* expression was observed in the lateral brain hemispheres and the pharynx, and a punctate pattern was observed throughout the dorsal side of the planarian but was conspicuously absent from the midline and the lateral body periphery. In contrast, *ephrin-2* expression was predominantly detected in the brain, ventral nerve cords, and pharynx. *ephrin-3* expression was detected in the dorsal head tip, brain, ventral nerve cords, pharynx, and in a punctate pattern throughout the body. Although we were intrigued by the expression of *ephrin-1* and hypothesized it might encode the ligand for *EphR1*, we proceeded to perform an agnostic screen for potential epistatic partners.

### *ephrin-1* RNAi phenocopies the *EphR1* knockdown phenotype

To investigate the role of candidate *Eph* and *ephrins* genes in planarian patterning, we screened and tested their function using RNAi and then processed animals for WISH to examine the patterns of mechanosensory neuron, intestinal, and CNS marker expression (Fig. 2 and Supplementary Fig. 5). We found that knockdown of *ephrin-1* resulted in ectopic expression of *pkd1L-2^+^* ciliated neurons, strikingly phenocopying *EphR1* knockdown (Fig. 2), strongly suggesting that *ephrin-1* and *EphR1* proteins may act as a ligand-receptor pair to facilitate mechanosensory neuron patterning. This phenotype was not observed for any other Ephrin signaling genes tested. In contrast, no obvious differences were observed in intestinal or central nervous system patterning, visualized with *cofilin* and *chat,* respectively, following knockdown of Ephrin signaling candidate genes (Supplementary Fig. 5). Taken together, these results suggest that Ephrin signaling does not globally regulate planarian tissue patterning but rather specific cell populations. The *ephrin-1* RNAi knockdown phenotype, together with complementary expression domains, is consistent with a ligand-receptor interaction that establishes a local exclusion zone for mechanosensory neuron differentiation or migration. This arrangement resembles Eph-ephrin signaling-mediated boundary sharpening in other organisms but appears to operate in adult tissue homeostasis. Future expression analysis using fluorescent in situ hybridization and higher-resolution imaging should help refine potential cell-interacting domains in planarian tissues.

**Figure 2:**
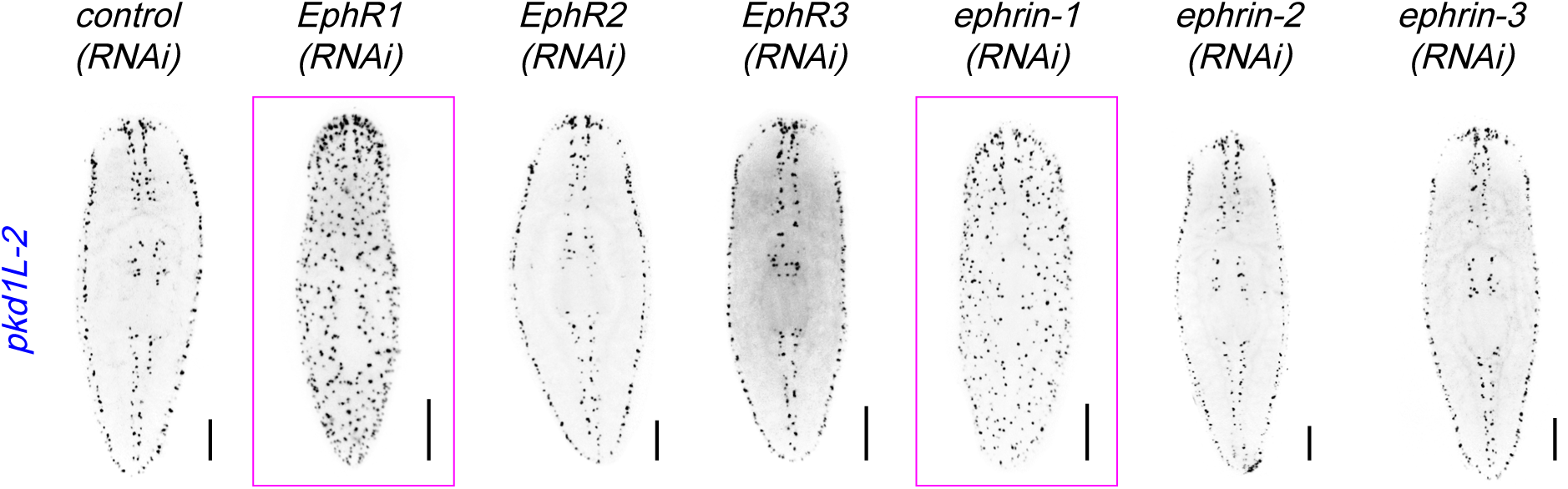
Effects of Ephrin signaling gene knockdown on peripheral sensory neuron expression. In situ hybridization for *pkd1L-2* was used to mark mechanosensory neurons (as shown in Fig. 1C) after RNAi of putative EphR and ephrin homologs. Knockdown of *ephrin-1* phenocopied *EphR1* knockdown. Animals are dorsal side up. Scale bars = 200 µm. Experiments were duplicated, with 5 worms per condition and 1 representative worm imaged.

Given the apparent relationship between *ephrin-1* and *EphR1*, *ephrin-1* was further investigated. The planarian single-cell database (Fincher et al., 2018) identified *ephrin-1* as enriched in the primary neural cluster and in subsets of neural cells, including ciliated and non-ciliated neurons, as well as in muscle cells. Although *ephrin-1* and *EphR1* are both expressed in the primary neural subcluster, their expression does not overlap (not shown). This finding was supported by WISH analysis of *EphR1* and *ephrin-1*. *ephrin-1* expression generally occurred in areas devoid of *EphR1* expression. Notably, ephrin-1 was not expressed along the dorsal ciliated stripe or the peripheral ciliated stripes. *ephrin-1* expression was also detected at the anterior pharynx and across the midline, regions that are also void of *EphR1* expression (Fig. 1A and Supplementary Fig. 2). Furthermore, the finding that *ephrin-1* is expressed in muscle suggests that the ligand may function as a position control gene to help regulate the planarian body plan (Reddien, 2018). Taken together, these data support a model in which *ephrin-1* and *EphR1* act in a common pathway to restrict the distribution and abundance of mechanosensory neurons. Because RNAi phenotypes can accumulate over time in planarians, we next examined the temporal emergence of the *EphR1* and *ephrin-1* knockdown phenotypes.

### EphR1–ephrin-1 signaling regulates sensory neuron patterning during adult tissue homeostasis

The striking *EphR1* and *ephrin-1* RNAi phenotype prompted us to investigate how their activity regulates the patterning of *pkd1L-2^+^* and *hmcn-1-L^+^*cells during adult tissue homeostasis. To this end, we performed an RNAi time course (outlined in Fig. 3A). Seven days after RNAi initiation, *EphR1(RNAi)* animals exhibited few ectopic *pkd1L-2^+^*and *hmcn-1-L^+^* cells, with ectopic cells increasing over time (Fig. 3B-C). In contrast, ectopic *pkd1L-2^+^* and *hmcn-1-L^+^*cells following ephrin-1 knockdown appeared more slowly; there were no obvious changes until 14 and 21 days after RNAi initiation for *pkd1L-2* and *hmcn-1-L*, respectively, but clear expansion of ectopic cells was observed at later timepoints (Fig. 3B-C). However, simultaneous knockdown of *EphR1* and *ephrin-1* caused robust ectopic expression of both marker genes (Fig. 3B-C). Cell marker genes appeared lateral to the dorsal ciliated stripe 7 days after the first double RNAi feeding, and expression increased across the entire dorsal side of the planarian over time. We hypothesize that Ephrin signaling provides continuous cues that specify and maintain the location of *pkd1L-2^+^*and *hmcn-1-L^+^* neurons, and that even a minor disruption of this signaling causes spatial disorganization. To investigate whether ectopic expression represents an increase in the total number of mechanosensory organ neurons, we quantified *pkd1L-2^+^* and *hmcn-1-L^+^* cells in control and RNAi-treated groups. We found a significant increase in *pkd1L-2^+^* and *hmcn-1-L^+^*neuronal populations over time compared to control worms (Fig. 3D). This suggests that EphR1 signaling is not only directly implicated in the boundary maintenance of *pkd1L-2^+^* and *hmcn-1-L^+^* neurons but also influences their population size. The progressive increase in *pkd1L-2^+^*and *hmcn-1-L^+^* neurons also suggests that EphR1-ephrin-1 signaling limits mechanosensory neuron output over time. This increase could reflect elevated progenitor differentiation, altered maintenance or survival, or derepression of marker expression in existing cells. Regardless of the mechanism, the data indicate that Ephrin signaling coordinates spatial restriction with population-level control in an adult sensory neuron organ system.

**Figure 3.**
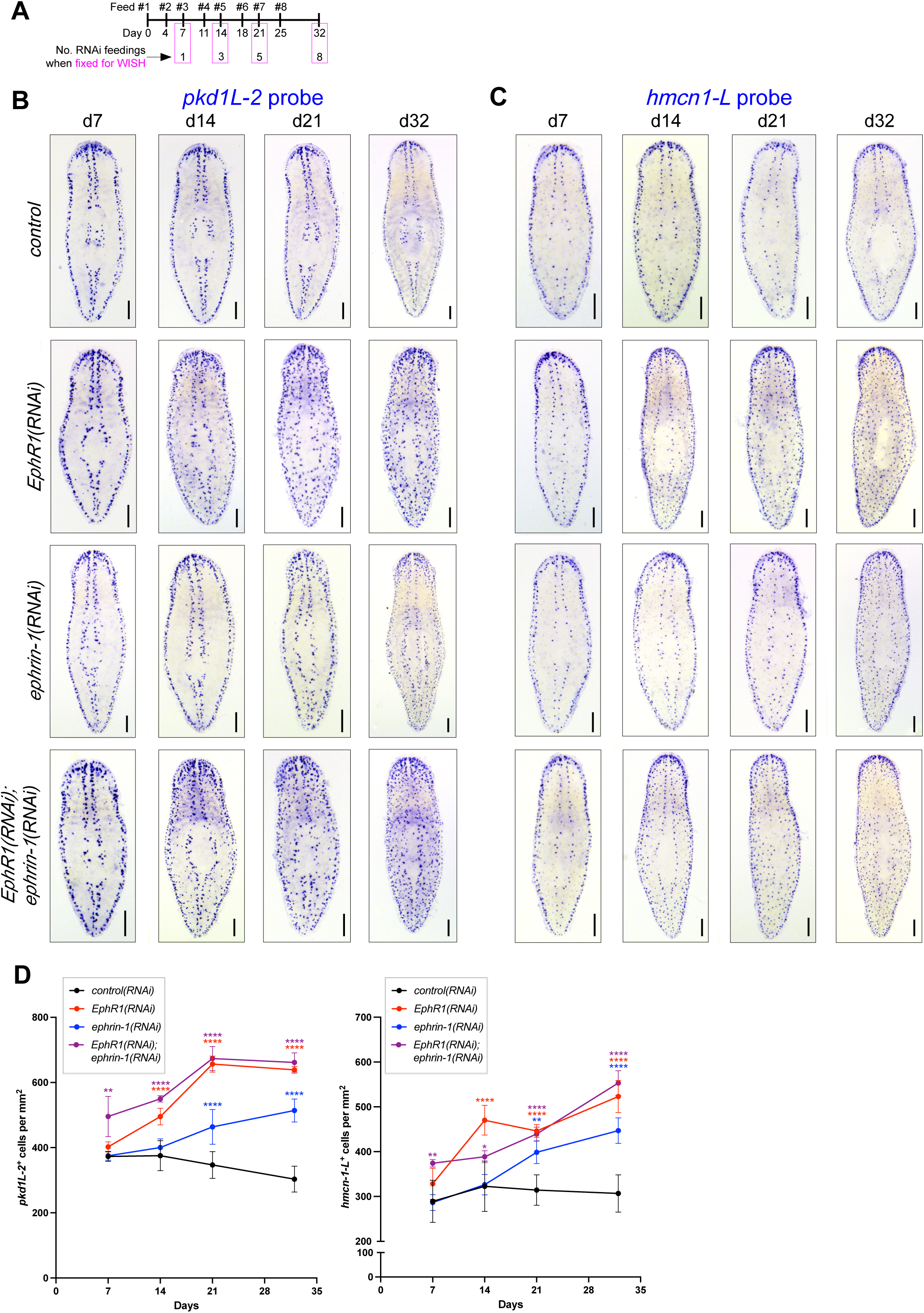
*EphR1* inhibition disrupts *pkd1L-2*^+^ and *hmcn-1-L*^+^ cell numbers over time. (A) RNAi time course and sample fixation scheme. (B-C) RNAi knockdown of *EphR1* disrupted the patterning of *pkd1L-2^+^* (B) and *hmcn-1-L^+^* (C) sensory neuron gene marker expression. Scale bars = 200 µm. Experiments were duplicated, with 3-5 worms per condition and 1 representative worm imaged. (D) Quantification of *pkd1L-2^+^* (B) and *hmcn-1-L^+^* (C) cells in control and RNAi samples. Statistical significance was assessed using two-way ANOVA with Dunnett’s multiple-comparisons test (*P < 0.05, **P < 0.01, ***P < 0.001, ****P < 0.0001).

### The *EphR1* RNAi ectopic sensory gene expression phenotype is stem cell-dependent

Quantitative analysis of the RNAi phenotypes strongly indicated that the animals are inappropriately generating additional mechanosensory cells. To test the hypothesis that the new cells arise from the neoblast population during normal cell turnover or as a result of disrupted signaling that feeds back to promote differentiation into more neurons, we performed irradiation assays. The planarian stem cell population is ablated within 24 hours when the animals are exposed to 100 Gy of ionizing radiation. We selected early time points based on our results (Figure 3). Animals treated with dsRNA were then subdivided, with one subgroup exposed to 100 Gy X-rays two days after the last RNAi feeding (Fig. 4A). All were processed for WISH to *pkdL1-2*. As we predicted, irradiated *EphR1(RNAi)* animals did not develop the ectopic expression phenotype, indicating that emergence of the ectopic population is neoblast-dependent (Fig. 4B).

**Figure 4.**
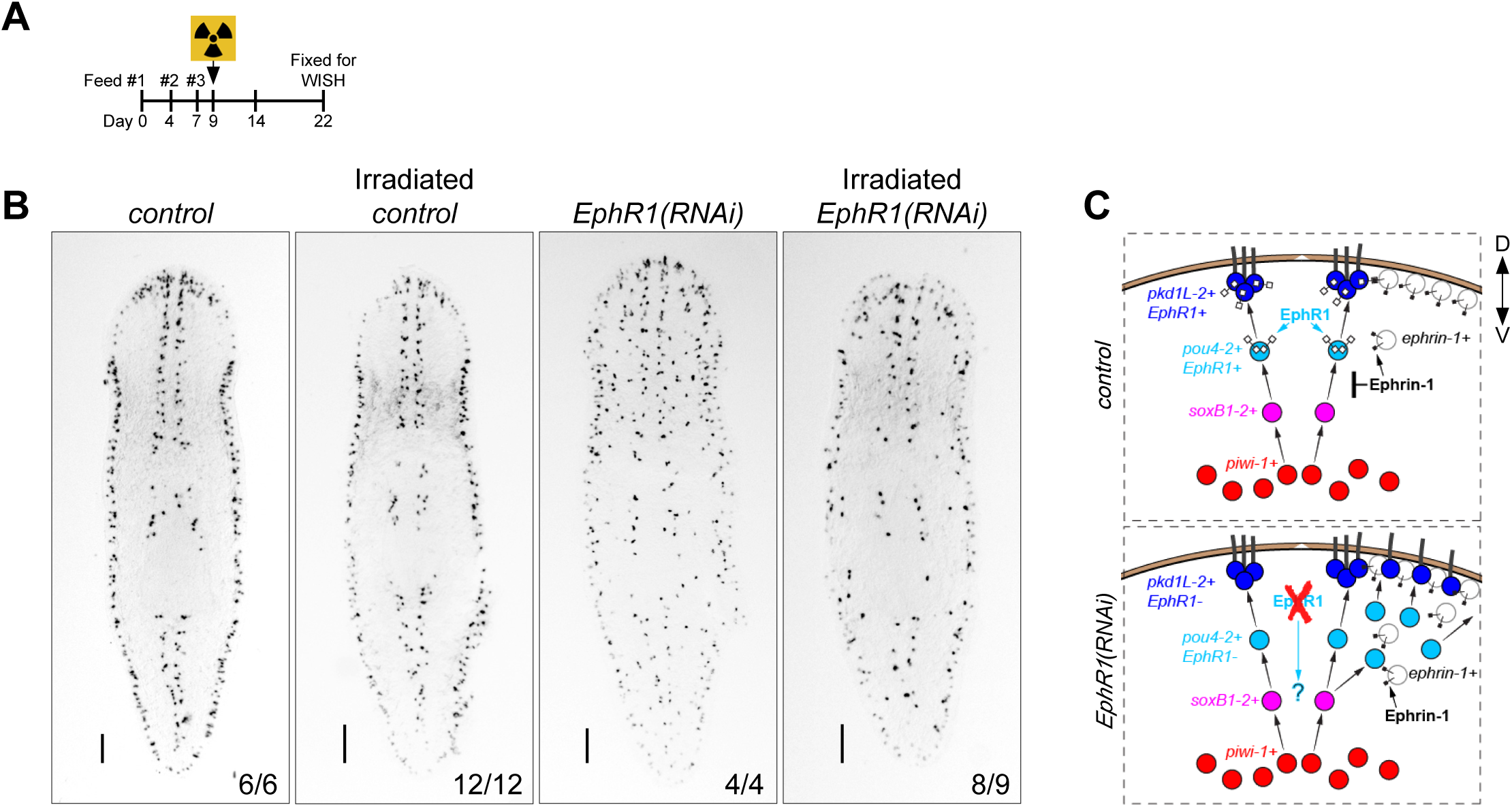
The *EphR1(RNAi)* ectopic sensory gene expression phenotype is stem cell-dependent. (A) RNAi time course, irradiation, and sample fixation scheme. (B) RNAi knockdown of *EphR1* disrupted the patterning of *pkd1L-2^+^* sensory neuron gene marker expression. Ectopic expression was abolished when planarians were treated with 100 Gy of X-rays two days after the third RNAi feeding, and animals were processed for WISH 13 days post-irradiation. Scale bars = 200 µm. (C) Working model for EphR1-ephrin-1 function in sensory neuron patterning.

To assess whether ectopic mechanosensory neurons alter gross mechanosensory responses, we tested tap-induced contraction (McCubbin et al., 2025; Ross et al., 2024; Ross et al., 2018) and observed no obvious loss of responsiveness in any RNAi condition (not shown). Higher-resolution assays will be required to determine whether ectopic cells integrate functionally and modify mechanosensory sensitivity.

Although no overt CNS changes were detected and we observed only visible expansion of the mechanosensory organ, we cannot rule out effects on other populations, as indicated by mispatterning defects in *cav1^+^* and *nkx2-4^+^* cells (Supplementary Fig. 1C). Other PNS populations not examined here might be affected by EphR1-ephrin-1 signaling. Future work is needed to comprehensively assess which cell types use EphR1-ephrin-1 signaling to establish and maintain patterning and proportionality. Furthermore, previous work showed that *soxB1-2* regulates *delta* expression, consistent with previous work implicating canonical Notch signaling in the generation and function of planarian sensory structures, including the differentiation of *pou4-2^+^* neurons (Elliott, 2016). Thus, future experiments could investigate the role of Notch signaling in the EphR1-ephrin-1 RNAi phenotype.

Notwithstanding, this research elucidates a role for Ephrin signaling in neural patterning in planarians. Using RNAi and WISH analysis, we showed that EphR1 is an essential regulator of a specific subset of peripheral sensory neurons. EphR1 signaling, mediated by interactions with ephrin-1-expressing cells, maintains the proper spatial positioning and population size of sensory neurons (Fig. 4C). Additionally, this research identified several Ephrin homologs with CNS-specific and other cell-type expression patterns, which should be useful for examining additional roles of Ephrin signaling in planarian tissue patterning and regeneration.

## Methods

### Planarian Culture

Asexual clonal line CIW4 of *S. mediterranea* was maintained in 1x Montjuïc salts in the dark at 20°C and fed weekly with pureed calf liver (Cebrià and Newmark, 2005; Merryman et al., 2018). Planarians 2-4 mm in length were starved one week prior to experimentation.

### Gene Identification and Cloning

The Pfam Ephrin receptor ligand binding domain (EPH_lbd) and Ephrin domain sequences (Mistry et al., 2021) were retrieved from the InterPro database (https://www.ebi.ac.uk/interpro/) and used to perform BLAST searches against the planarian dd_Smed_v6 transcriptome and the d_Smed_v6.PCFL.proteins proteome (Rozanski et al., 2019). We then extracted the predicted EPH_lbd and Ephrin domains from planarian hits and performed BLAST analysis. Together, these searches identified four Ephrin receptor and four ephrin genes; reciprocal BLAST and domain searches using SMART (Ponting et al., 1999) confirmed hits with confidence. The transcripts and annotations generated in Blast2GO (Gotz et al., 2008) and SMART are listed in Supplementary Table 3. Genes were obtained from an EST library (Zayas et al., 2005), cloned using gene-specific primers, or synthesized as eBlocks (IDT) and inserted into pPR-T4P plasmid vectors through ligation-independent cloning (see Supplementary Table 4). dd_Smed_v6 transcripts are referred to as *dd_transcript no.* for brevity.

### Phylogenetic analysis

Ephrin receptor and ligand homologs from a number of invertebrate and vertebrate species were adopted from (Mellott and Burke, 2008). The *Schmidtea mediterranea* nucleotide sequences were translated into amino acid sequences using the ExPASy Translate tool (Gasteiger et al., 2003) (Supplementary File 1). The software Geneious (www.geneious.com) was used to align the predicted proteins to the alignment files from Mellott and Burke (2008) using the Clustal Omega plugin. Protein alignments were manually inspected prior to performing Bayesian inference of phylogeny using the MrBayes 3.2.6 (Huelsenbeck and Ronquist, 2001) plugin developed by Marc Suchard and the Geneious Team with the following settings: unconstrained branch length, shape parameter exponential of 10, 1.1 million chain length, 4 heated chains, 0.2 heated chain temp., WAG substitution model, gamma rate variation model, 10% burnin length with subsampling frequency of 200. The protein alignments are provided in Supplementary Files S2-9.

### RNA Interference

Double-stranded RNA was prepared via bacterial induction as described in Adler and Alvarado (2018). Briefly, plasmids containing target genes were introduced into HT115-competent cells that can be induced to express T7 polymerase in the presence of isopropyl β-d-thiogalactoside (IPTG). Overnight cultures were diluted into 2xYT media and incubated for 2 hours at 37°C with shaking. Upon reaching an OD_600_ of approximately 0.6-1.0, cultures were induced with 0.4 mM IPTG and incubated for an additional 2 hours at 37°C with shaking. dsRNA was pelleted and stored at -80°C. For RNAi feeding, dsRNA was mixed with liver; *gfp* dsRNA was used as a control. Two RNAi paradigms were used in this study. For endpoint analyses, animals were fed dsRNA 8 times over 4 weeks and were fixed 7 days after the final feeding. For RNAi time-course experiments, animals received repeated dsRNA feedings according to the schedule shown, and samples were collected at defined time points relative to the first feeding (Fig 3A and 4A).

### *In Situ* Hybridization

Riboprobes were synthesized using an in vitro transcription reaction with a DNA template and digoxigenin-labeled NTPs (Pearson et al., 2009). Whole-mount *in-situ* hybridization was performed as previously described (King and Newmark, 2013). Briefly, animals were sacrificed in 5% N-acetyl-cysteine solution for 5 minutes and fixed in 4% formaldehyde solution for 20 minutes. Samples were bleached for 4 hours in a formamide-based solution, permeabilized in proteinase K solution, and post-fixed in a 4% formaldehyde solution. Probe hybridization, antibody incubation, and wash steps were performed using an InsituPro VS liquid-handling robot (CEM). Samples were incubated with anti-Digoxigenin-AP (1:2000, Roche) for chromogenic detection, and the signals were subsequently developed with NBT/BCIP in AP buffer.

### Immunostaining

The protocol was carried out as previously described (Ross et al., 2015). Animals were sacrificed in ice-cold 2% HCl for 5 minutes, followed by incubation in Carnoy’s fixative (6 parts ethanol: 3 parts CHCl_3_: 1 part glacial acetic acid) for 2 hours at 4°C. Animals were dehydrated with 100% methanol for 1 hour at 4°C, then bleached overnight under a lamp with 6% H_2_O_2_ in methanol. Animals were then rehydrated in 75%, 50%, and 25% methanol-PBSTx, followed by two 5-minute PBSTx washes. PBSTb (1% BSA in PBSTx) was used for blocking at room temperature for 4 hours. Primary antibody labeling was performed with 3C11 (anti-Synapsin-1), and the samples were incubated in the dark for 16 hours at 4°C. Six 1-hour PBSTx washes, followed by 1 hour of PBSTb blocking, were performed prior to incubation with anti-mouse-HRP (1:1000, Cell Signaling) for 16 hours in the dark at 4°C. After six 1-hour PBSTx washes, Synapsin was detected through TSA development. Briefly, animals were incubated in borate buffer for 5 minutes, then TSA reaction buffer (1:250 Cy3-tyramide) for 10 minutes. Additional H_2_O_2_ was spiked in, and animals were incubated for 10 more minutes.

### X-ray irradiation

RNAi-treated animals were irradiated with 100 Gy of X-rays (130 kV, 5 mA, 8.4 Gy/min) for approximately 12 min using a CellRad irradiator (Precision X-Ray, Madison, CT).

### Imaging

Animals processed for *in situ* hybridization were imaged using a Leica DFC450 camera mounted on a Leica M205 stereoscope. Animals processed for immunohistochemistry were mounted in Vectashield diluted in 80% glycerol and imaged using a Zeiss AxioZoom v.16 equipped with an Apotome via ZEN Pro software.

### Cell Counting and Quantification

Counts of *pkd1L-2* and *hmcn-1-L* cells were performed using ImageJ v1.53. Cell quantification was performed automatically using the particle analysis function on ImageJ. Graphs were made using GraphPad Prism. Statistical significance was assessed using a two-way ANOVA followed by Dunnett’s multiple comparisons test. Differences were considered significant at P < 0.05.

## Acknowledgements

We thank Mallory Cathell, Deborah Yelon, and members of the Zayas Lab for helpful discussions about this work. The anti-SYNORF1 (3C11) antibodies were obtained from the Developmental Studies Hybridoma Bank, created by the Eunice Kennedy Shriver National Institute of Child Health and Human Development (NICHD) of the NIH and maintained at The University of Iowa, Department of Biology, Iowa City, IA 52242. This work was supported by a California Institute for Regenerative Medicine (CIRM) postdoctoral fellowship (EDUC4-12813) to M.A.A.; SDB Choose Development! funding to Christian Torres; and NIH R01GM135657 to R.M.Z.

## Author contributions

K.G.R., R.A.M. and R.M.Z. conceived the project; K.G.R. and R.M.Z. supervised the research; M.A.A., S.W., R.A.M., A.L.F. A.M., J.M.S., C.T., R.M.Z. performed experiments and analyzed data; M.A.A., S.W., R.A.M, K.G.R., and R.M.Z. interpreted the results, prepared the figures and illustrations, and wrote the manuscript.

## Supplemental Figure Legends

**Supplemental Figure 1:**
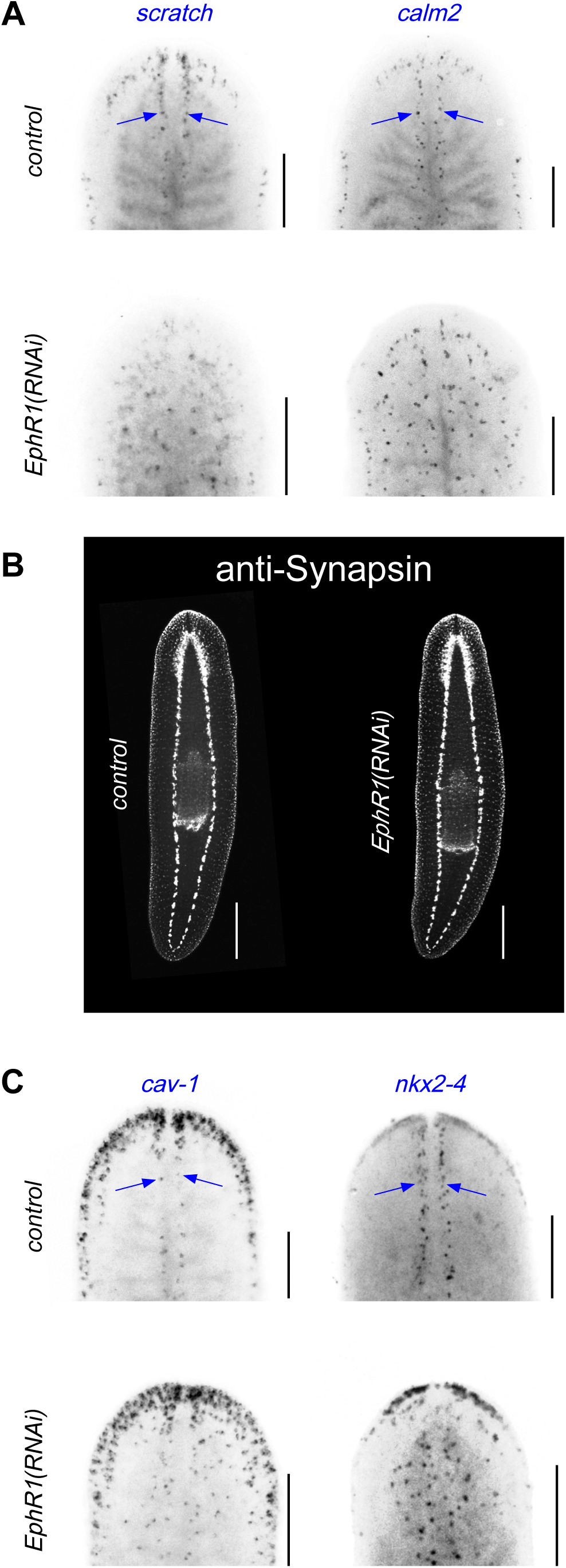
Analysis of genes expressed in *EphR1^+^*single-cell RNA-seq clusters. (A) Knockdown of *EphR1* resulted in ectopic expression of *scratch*^+^ and calm2+. (B) CNS architecture revealed by anti-Synapsin staining. Scale bars = 200 µm. (C) Knockdown of *EphR1* resulted in ectopic expression of *cav-1*^+^ and *nkx2-4*^+^ cells. Scale bars = 200 µm. All animals are dorsal side up. Experiments were duplicated, with 5 worms per condition and 1 representative worm imaged per condition.

**Supplemental Figure 2.**
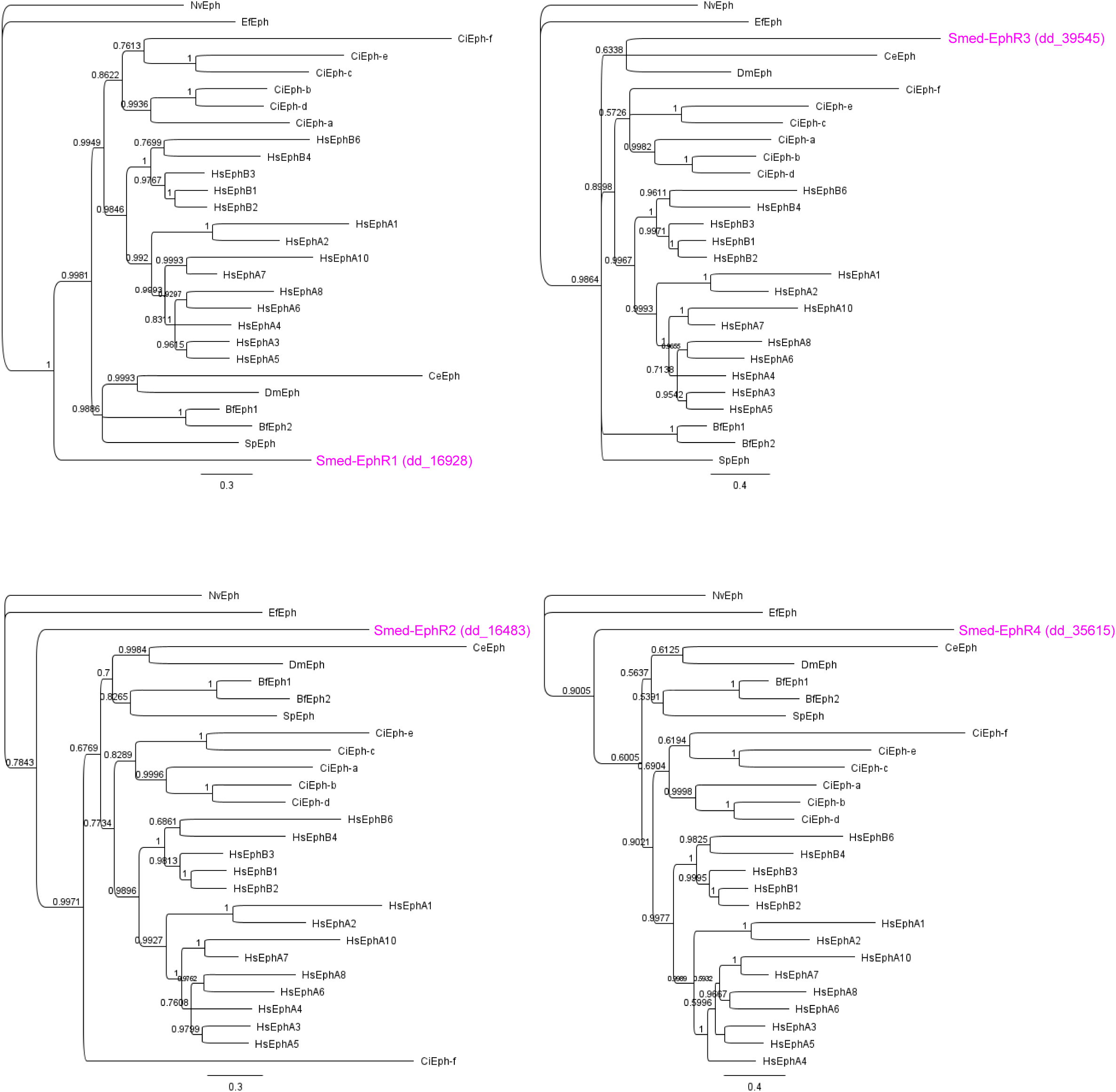
A Bayesian inference phylogeny of Ephrin Receptor proteins. Node support values are shown as percentages next to the relevant nodes. *Bf, Brachiostoma floridae; Ce, Caenorhabditis elegans; Ci, Ciona intestinalis; Dm, Drosophila melanogaster; Efn, Ephydatia fluviatilis; Hs, Homo sapiens; Nv, Nematostella vectensis; Sp, Strongylocentrotus purpuratus; Smed, Schmidtea mediterranea*.

**Supplemental Figure 3.**
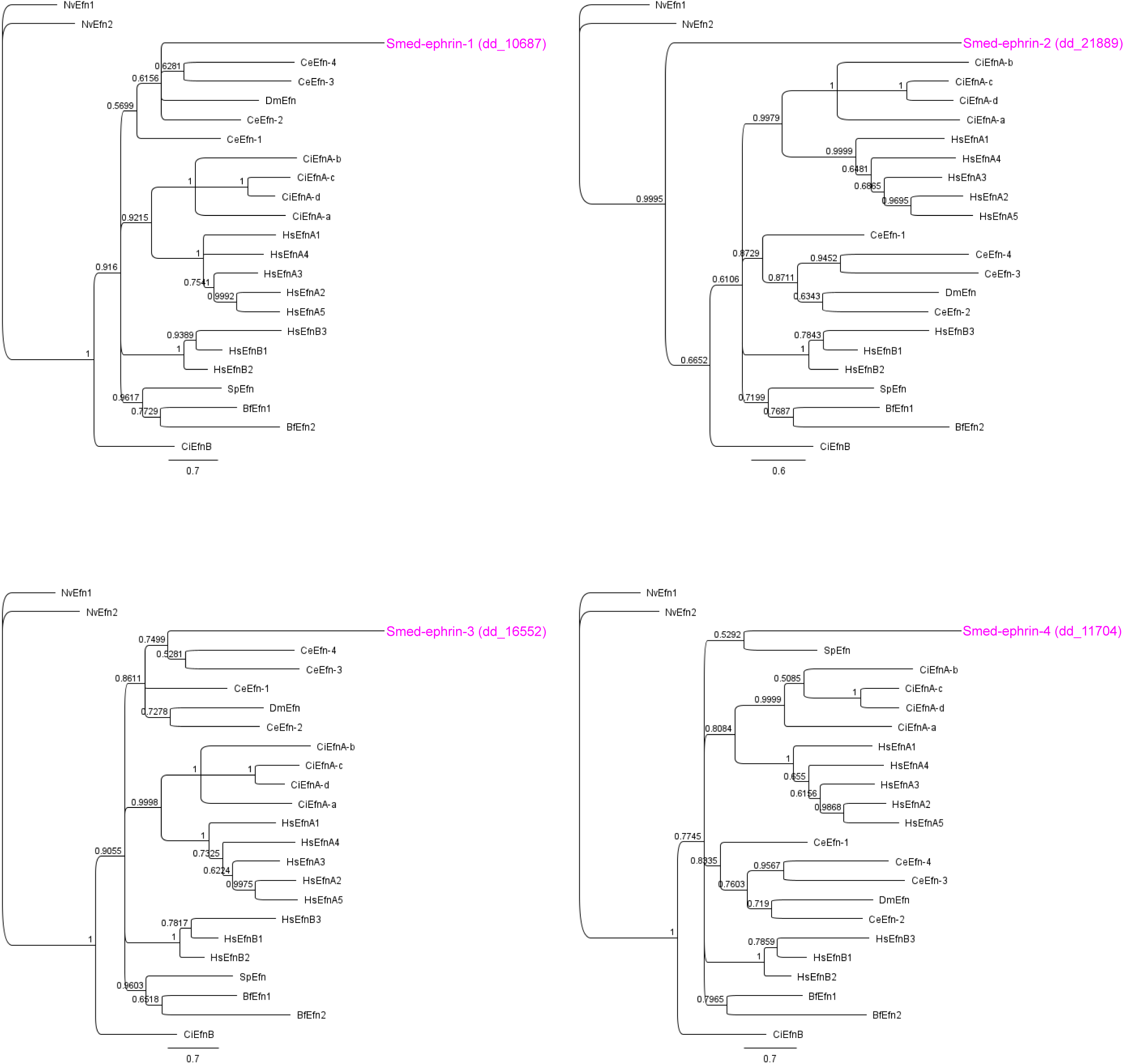
A Bayesian phylogenetic tree of ephrin ligand proteins. Node support values are shown as percentages next to the relevant nodes. *Bf, Brachiostoma floridae; Ce, Caenorhabditis elegans; Ci, Ciona intestinalis; Dm, Drosophila melanogaster; Efn, Ephydatia fluviatilis; Hs, Homo sapiens; Nv, Nematostella vectensis; Sp, Strongylocentrotus purpuratus; Smed, Schmidtea mediterranea*.

**Supplemental Figure 4.**
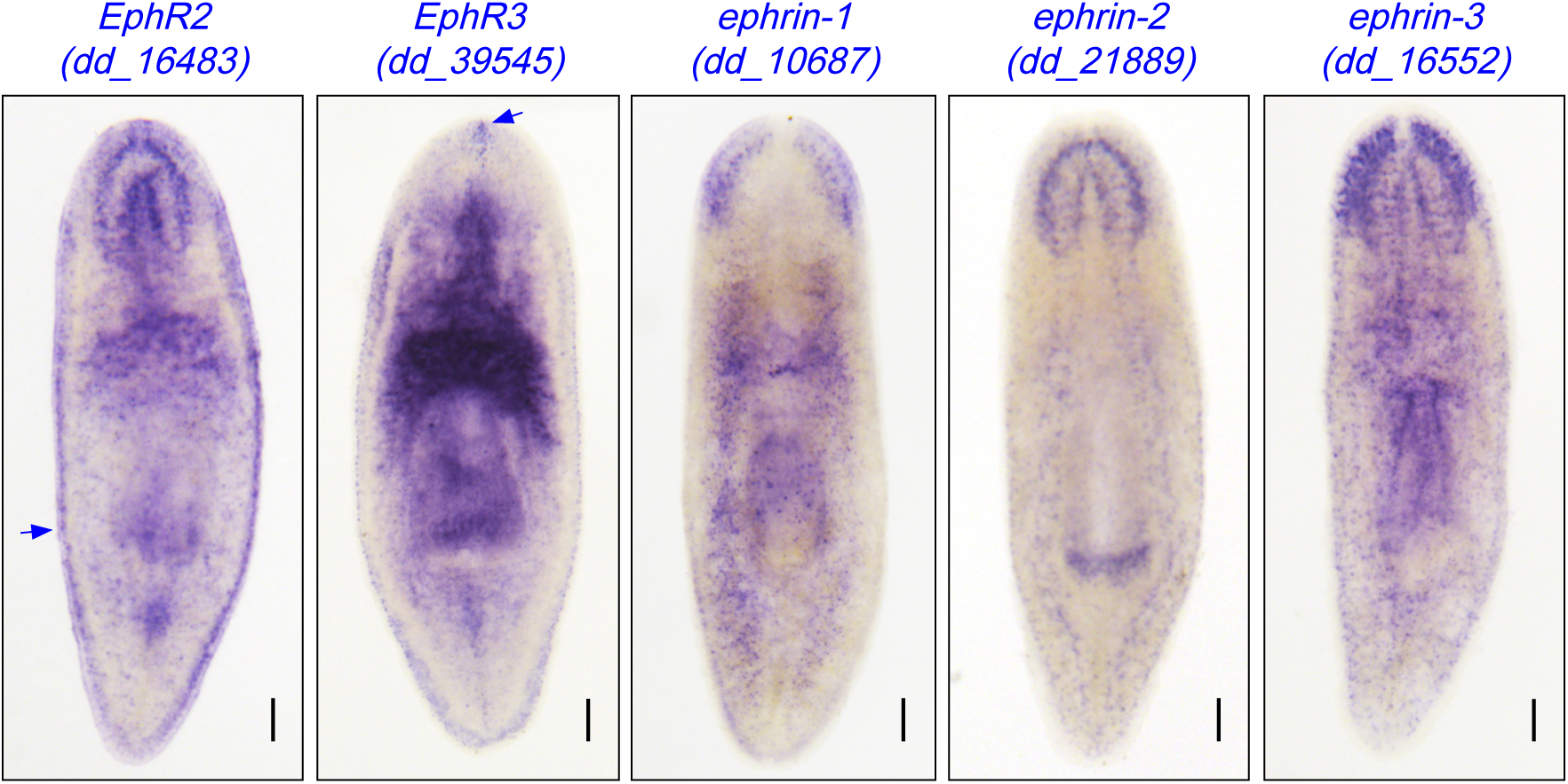
Expression patterns of candidate EphRs and ephrin genes. WISH analysis shows expression patterns of the two Eph receptors (EphRs) and three ephrin ligand genes examined to date. Scale bars = 200 µm.

**Supplemental Figure 5.**
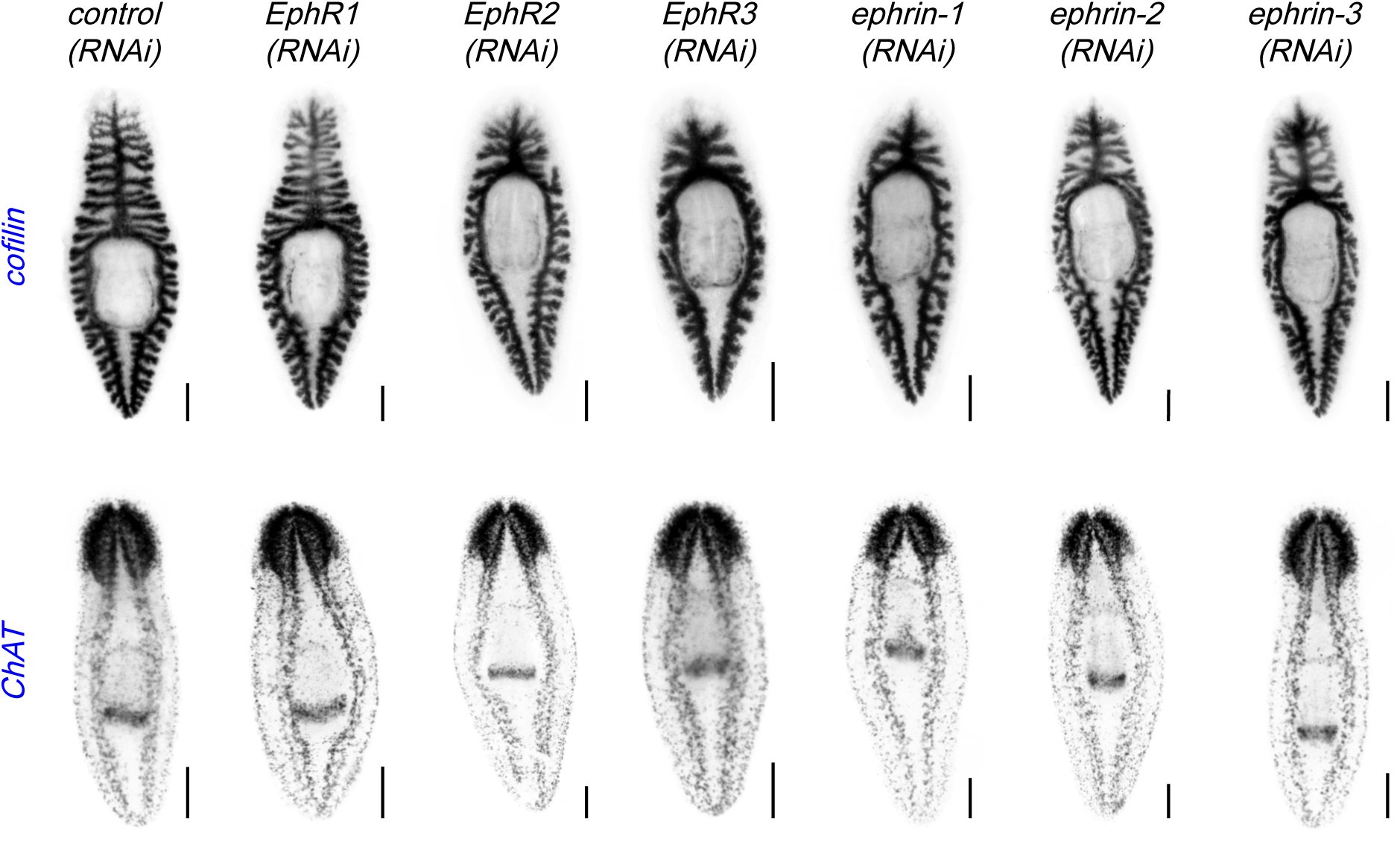
Effects of ephrin signaling gene knockdown on peripheral sensory neuron expression and on intestinal and central nervous system architecture. In situ hybridization for *cofilin* and *chat* was used to mark the intestine and the nervous system, respectively. Knockdown of ephrin signaling genes did not result in obvious defects in intestinal or CNS morphology. Scale bars = 200 µm. Experiments were duplicated, with 5 worms per condition and 1 representative worm imaged.

## References

Aberle, H. (2019). Axon Guidance and Collective Cell Migration by Substrate-Derived Attractants. Frontiers in Molecular Neuroscience 12.

Adler, C. E. and Alvarado, A. S. (2018). Systemic RNA Interference in Planarians by Feeding of dsRNA Containing Bacteria. Methods Mol Biol 1774, 445–454.

Arcas, A., Wilkinson, D. G., Nieto, M. Á. and Rogers, R. (2020). The Evolutionary History of Ephs and Ephrins: Toward Multicellular Organisms. Molecular biology and evolution. 37, 379–394.

Cebrià, F. (2007). Regenerating the central nervous system: how easy for planarians! Dev Genes Evol 217, 733–748.

Cebrià, F. (2008). Organization of the nervous system in the model planarian Schmidtea mediterranea: an immunocytochemical study. Neurosci Res 61, 375–384.

Cebrià, F. and Newmark, P. A. (2005). Planarian homologs of netrin and netrin receptor are required for proper regeneration of the central nervous system and the maintenance of nervous system architecture. Development 132, 3691–3703.

Elliott, S. A. (2016). Studies of conserved cell-cell signaling pathways in the planarian, Schmidtea mediterranea. In Department of Neurobiology and Anatomy: University of Utah.

Fincher, C. T., Wurtzel, O., de Hoog, T., Kravarik, K. M. and Reddien, P. W. (2018). Cell type transcriptome atlas for the planarian Schmidtea mediterranea. Science 360, eaaq1736.

Gasteiger, E., Gattiker, A., Hoogland, C., Ivanyi, I., Appel, R. D. and Bairoch, A. (2003). ExPASy: The proteomics server for in-depth protein knowledge and analysis. Nucleic Acids Res 31, 3784–3788.

Goldshmit, Y., Galea, M. P., Wise, G., Bartlett, P. F. and Turnley, A. M. (2004). Axonal Regeneration and Lack of Astrocytic Gliosis in EphA4-Deficient Mice. Journal of Neuroscience 24, 10064–10073.

Gotz, S., Garcia-Gomez, J. M., Terol, J., Williams, T. D., Nagaraj, S. H., Nueda, M. J., Robles, M., Talon, M., Dopazo, J. and Conesa, A. (2008). High-throughput functional annotation and data mining with the Blast2GO suite. Nucleic Acids Res 36, 3420–3435.

Huelsenbeck, J. P. and Ronquist, F. (2001). MRBAYES: Bayesian inference of phylogenetic trees. Bioinformatics 17, 754–755.

Ivankovic, M., Haneckova, R., Thommen, A., Grohme, M. A., Vila-Farre, M., Werner, S. and Rink, J. C. (2019). Model systems for regeneration: planarians. Development 146.

Jiao, J.-w., Feldheim, D. A. and Chen, D. F. (2008). Ephrins as negative regulators of adult neurogenesis in diverse regions of the central nervous system. Proceedings of the National Academy of Sciences 105, 8778–8783.

Kandel, E., Schwartz, J., Jessell, T., Siegelbaum, S., Hudspeth, A. and Mack, S. (2014). Patterning the Nervous System (5th Edition edn). New York, NY: McGraw-Hill Education.

Kania, A. and Klein, R. (2016). Mechanisms of ephrin-Eph signalling in development, physiology and disease. Nat Rev Mol Cell Biol 17, 240–256.

King, H. O., Owusu-Boaitey, K. E., Fincher, C. T. and Reddien, P. W. (2024). A transcription factor atlas of stem cell fate in planarians. Cell Reports 43, 113843.

King, R. S. and Newmark, P. A. (2013). In situ hybridization protocol for enhanced detection of gene expression in the planarian Schmidtea mediterranea. BMC Developmental Biology 13, 8.

Klein, R. (2012). Eph/ephrin signalling during development. Development 139, 4105–4109.

Laussu, J., Khuong A Fau - Gautrais, J., Gautrais J Fau - Davy, A. and Davy, A. (2014). Beyond boundaries--Eph:ephrin signaling in neurogenesis. Cell Adh Migr 8, 345–359.

Lisabeth, E. M., Falivelli, G. and Pasquale, E. B. (2013). Eph receptor signaling and ephrins. Cold Spring Harb Perspect Biol 5.

McCubbin, R. (2022). The role of pou4-2 in mechanosensory neuron function. In Biology. [unpublished Master’s thesis]: San Diego State University.

McCubbin, R. A., Auwal, M. A., Wang, S., Alvarez Zepeda, S., Sasik, R., Zeller, R. W., Ross, K. G. and Zayas, R. M. (2025). Smed-pou4-2 regulates mechanosensory neuron regeneration and function in planarians. Elife 14.

Mellott, D. O. and Burke, R. D. (2008). The molecular phylogeny of eph receptors and ephrin ligands. BMC Cell Biol 9, 27.

Merryman, M. S., Alvarado, A. S. and Jenkin, J. C. (2018). Culturing Planarians in the Laboratory. Methods in molecular biology 1774, 241–258.

Mistry, J., Chuguransky, S., Williams, L., Qureshi, M., Salazar, G. A., Sonnhammer, E. L. L., Tosatto, S. C. E., Paladin, L., Raj, S., Richardson, L. J., et al. (2021). Pfam: The protein families database in 2021. Nucleic Acids Res 49, D412–D419.

Newmark, P. A. and Sánchez Alvarado, A. (2002). Not your father’s planarian: a classic model enters the era of functional genomics. Nat Rev Genet 3, 210–219.

Pearson, B. J., Eisenhoffer, G. T., Gurley, K. A., Rink, J. C., Miller, D. E. and Sánchez Alvarado, A. (2009). Formaldehyde-based whole-mount in situ hybridization method for planarians. Developmental Dynamics 238, 443–450.

Petersen, C. P. and Reddien, P. W. (2009). A wound-induced Wnt expression program controls planarian regeneration polarity. PNAS 106, 17061 – 17066.

Ponting, C. P., Schultz, J., Milpetz, F. and Bork, P. (1999). SMART: identification and annotation of domains from signalling and extracellular protein sequences. Nucleic Acids Res 27, 229–232.

Reddien, P. W. (2018). The Cellular and Molecular Basis for Planarian Regeneration. Cell 175, 327–345.

Reddien, P. W. (2022). Positional Information and Stem Cells Combine to Result in Planarian Regeneration. Cold Spring Harb Perspect Biol 14, a040717.

Reddien, P. W. and Sánchez Alvarado, A. (2004). Fundamentals of planarian regeneration. Annu Rev Cell Dev Biol 20, 725–757.

Ross, K. G., Alvarez Zepeda, S., Auwal, M. A., Garces, A. K., Roman, S. and Zayas, R. M. (2024). The Role of Polycystic Kidney Disease-Like Homologs in Planarian Nervous System Regeneration and Function. Integr Org Biol 6, obae035.

Ross, K. G., Currie, K. W., Pearson, B. J. and Zayas, R. M. (2017). Nervous system development and regeneration in freshwater planarians. Wiley Interdiscip Rev Dev Biol 6.

Ross, K. G., Molinaro, A. M., Romero, C., Dockter, B., Cable, K. L., Gonzalez, K., Zhang, S., Collins, E. S., Pearson, B. J. and Zayas, R. M. (2018). SoxB1 Activity Regulates Sensory Neuron Regeneration, Maintenance, and Function in Planarians. Dev Cell 47, 331–347.e335.

Ross, K. G., Omuro, K. C., Taylor, M. R., Munday, R. K., Hubert, A., King, R. S. and Zayas, R. M. (2015). Novel monoclonal antibodies to study tissue regeneration in planarians. BMC Developmental Biology 15, 2.

Rozanski, A., Moon, H., Brandl, H., Martin-Duran, J. M., Grohme, M. A., Huttner, K., Bartscherer, K., Henry, I. and Rink, J. C. (2019). PlanMine 3.0-improvements to a mineable resource of flatworm biology and biodiversity. Nucleic Acids Res 47, D812–D820.

Wang, S. (2019). The role of POU4 homologue in the regeneration and maintenance of planarian sensory neurons. In Biology: San Diego State University.

Wilkinson, D. G. (2014). Regulation of cell differentiation by Eph receptor and ephrin signaling. Cell Adh Migr 8, 339–348.

Yang, J. S., Wei, H. X., Chen, P. P. and Wu, G. (2018). Roles of Eph/ephrin bidirectional signaling in central nervous system injury and recovery. Exp Ther Med 15, 2219–2227.

Yazawa, S., Umesono, Y., Hayashi, T., Tarui, H. and Agata, K. (2009). Planarian Hedgehog/Patched establishes anterior–posterior polarity by regulating Wnt signaling. PNAS 106, 22329–22334.

Zayas, R. M., Hernandez, A., Habermann, B., Wang, Y., Stary, J. M. and Newmark, P. A. (2005). The planarian Schmidtea mediterranea as a model for epigenetic germ cell specification: analysis of ESTs from the hermaphroditic strain. Proceedings of the National Academy of Sciences of the United States of America 102, 18491–18496.

